# Lost in space(s): multimodal neuroimaging of disorientation along the Alzheimer’s disease continuum

**DOI:** 10.1101/2023.01.25.525587

**Authors:** Gregory Peters-Founshtein, Lidor Gazit, Tahel Naveh, Liran Domachevsky, Amos Korczyn, Hanna Bernstine, David Groshar, Gad A. Marshall, Shahar Arzy

**Author notes:** **Correspondence should be addressed to:** Shahar Arzy, MD PhD, Department of Medical Neurobiology, Faculty of Medicine, Hebrew University of Jerusalem, Jerusalem, Israel.

## Abstract

Orientation is a fundamental cognitive faculty, allowing the behaving self to link his/her current state to their internal representations of the external world. Once exclusively linked to knowledge of the current place and present time, in recent years, the concept of orientation has evolved to include processing of social, temporal, and abstract relations. Concordantly with the growing focus on orientation, spatial disorientation has been increasingly recognized as a hallmark symptom of Alzheimer’s disease (AD). However, few studies have sought to explore disorientation along the AD continuum beyond the spatial domain.

51 participants along the AD continuum performed an orientation task in the spatial, temporal and social domains. Under functional magnetic resonance imaging (fMRI), participants determined which of two familiar places/events/people is geographically/chronologically/socially closer to them, respectively. A series of analyses revealed disorientation along the AD-continuum to follow a three-way association between (1) orientation domain, (2) brain region, and (3) disease stage. Specifically, participants with MCI exhibited impaired spatio-temporal orientation and reduced task-evoked activity in temporoparietal regions, while participants with AD dementia exhibited impaired social orientation and reduced task-evoked activity in frontoparietal regions. Furthermore, these patterns of hypoactivation coincided with Default Mode Network (DMN) sub-networks, with spatio-temporal orientation activation overlapping DMN-C and social orientation with DMN-A. Finally, these patterns of disorientation-associated hypoactivations coincided with patterns of fluorodeoxyglucose (FDG) hypometabolism and cortical atrophy characteristic to AD-dementia.

Taken together, our results suggest that AD may constitute a disorder of orientation, characterized by a biphasic process as (1) early spatio-temporal and (2) late social disorientation, concurrently manifesting in task-evoked and neurodegenerative changes in temporoparietal and parieto-frontal brain networks, respectively. We propose that a profile of disorientation across multiple domains offers a unique window into the progression of AD.

## Introduction

Orientation is a fundamental cognitive faculty, allowing the behaving self to link his/her current state to their internal representations of the external world (Berrios, 1982; Peer, Salomon, Goldberg, Blanke, & Arzy, 2015). Commonly, orientation is evaluated in the spatial, temporal and social domains, and as such it is recognized as the bedrock of the neurological clinical evaluation (Mahendran, Chua, Feng, Kua, & Preedy, 2015; Rapoport & Rapoport, 2015). Nonetheless, standard evaluations of orientation are limited to testing only the patient’s knowledge about the present time, current location and personal identity, resulting in low sensitivity to early cognitive decline (Peters-founshtein et al., 2018).

In recent years, several lines of research (Coughlan, Laczó, Hort, Minihane, & Hornberger, 2018; DeIpolyi, A. R., Rankin, K. P., Mucke, L., Miller, B. L., Gorno-Tempini, 2007; El Haj & Antoine, 2018; Kunz et al., 2015; Peters-Founshtein et al., 2018) have demonstrated that spatial orientation is potentially affected early on by AD pathology. One such study (Coughlan et al., 2019) used the Sea Hero Quest (SHQ) spatial navigation paradigm to compare young, cognitively intact, heterozygote carriers of Apolipoprotein E (APOE)-ε4 alleles (ε3/ε4), a known risk-multiplier of AD to demographically-matched healthy homozygote (ε3/ε3) participants. Comparing the two groups as they perform several goal-oriented wayfinding tasks revealed significant disruptions in navigation performance in people at-risk for AD showing no clinically detectable cognitive deficits. However, orientation is not restricted to the spatial domain. It involves other domains such as the temporal and social ones (Du, Basyouni, & Parkinson, 2021; Parkinson, Liu, & Wheatley, 2014; Peer et al., 2015), that have been shown to be progressively impaired along the AD-continuum (Dafni-Merom, Peters-Founshtein, Kahana-Merhavi, & Arzy, 2019; Peters-Founshtein et al., 2018). Moreover, tests of orientation have been found to better discriminate between cognitively normal (CN) and mild cognitive impairment (MCI) participants (95% accuracy) when compared to standard neuropsychological evaluations (Addenbrooke’s Cognitive Examination (ACE) – 71%, Mini Mental State Examination (MMSE) – 70%) (Peters-Founshtein et al., 2018). This superiority may stem from a considerable overlap between the patterns of orientation-evoked brain activity and patterns of AD neurodegeneration (Peters-Founshtein et al., 2018).

Independently, the pattern of orientation-evoked brain activity was found to markedly overlap with the Default Mode Network (DMN) (Hayman & Arzy, 2021; Peer et al., 2015; Peters-Founshtein et al., 2018). The DMN is a network of interconnected brain regions, active when individuals engage in self-referential tasks such as autobiographical memory retrieval and future planning (Buckner, Andrews-Hanna, & Schacter, 2008). Furthermore, AD neuropathology (amyloid-β (Aβ) and tau) has been shown to carry increased probability of spreading within rather than outside of the DMN (Adams, Maass, Harrison, Baker, & Jagust, 2019; Buckner et al., 2005; Franzmeier, Dewenter, et al., 2020; Franzmeier, Neitzel, et al., 2020). Subsequent studies, re-evaluating DMN homogeneity, have suggested the DMN is comprised of partially dissociated components, each underlying different cognitive functions (Andrews-Hanna, Reidler, Sepulcre, Poulin, & Buckner, 2010; A. J. Barnett et al., 2020, 2021; D. A. Barnett, Arnold, Valenzuela, Brayne, & Schneider, 2014; Buckner & DiNicola, 2019). Taken together, the overlapping patterns of DMN connectivity and orientation in space, time, and person, imply a latent model of inter-related neuropathological and cognitive changes in AD (Buckner et al., 2005).

In the current comprehensive study, we aimed to evaluate the relations between (1) spatio-temporal and social disorientation, (2) DMN subnetworks and (3) neurodegeneration, in individuals along the AD continuum, using positron emission tomography (PET)-functional magnetic resonance imaging (fMRI). Considering the neuropsychological profile of AD-related cognitive decline (early spatio-temporal and later social decline), jointly with multiple studies suggesting a segregation between spatio-temporal and social processing in the brain, we hypothesized that spatio-temporal and social orientation would be differently affected along the AD-continuum. Hence, we set to test and characterize these differences in behavioral performance and brain activity in the context of DMN topology and patterns of AD-related neurodegeneration.

## Methods

### Participants

Fifty-one individuals (27 females, mean age 71.43±0.82, for detailed demographics see Table 1) participated in the study, including 35 cognitively impaired participants (12 with AD dementia and 23 with amnestic MCI) and 16 age-matched CN older adults. Participants underwent a complete neurological examination, cerebrovascular risk-factor assessment using the Hachinski Ischemic Scale (Hachinski et al., 1975), and a comprehensive neuropsychological evaluation that included the Clinical Dementia Rating (Morris, 1993), Montreal Cognitive Assessment (Nasreddine et al., 2005), ACE (Mathuranath, Nestor, Berrios, Rakowicz, & Hodges, 2000), and Frontal Assessment Battery (Dubois, Slachevsky, Litvan, & Pillon, 2000). Cognitively impaired participants met the National Institute on Aging and the Alzheimer’s Association clinical criteria for AD-dementia or amnestic-MCI (Albert et al., 2011; S. Gauthier et al., 2006; Mckhann et al., n.d.; Petersen et al., 1999). All participants underwent structural T1 and T2 weighted MRI and fluorodeoxyglucose (FDG)-PET, which were reviewed by neuroradiology and nuclear medicine specialists to exclude non-AD etiologies. All participants provided written informed consent prior to undergoing study procedures, and the study was approved by the ethics committee of the Assuta Medical Center.

**Table 1.** Demographics and neuropsychological assessment scores.

|  | CN | MCI | AD dementia |
| --- | --- | --- | --- |
| Gender (F \| M) | 9 \| 7 | 14 \| 9 | 4 \| 8 |
| Age (years) | 69.5 ± 1.2 | 73.13 ± 1.38 | 70.75 ± 1.91 |
| Education (years) | 17.31 ± 0.8 | 16.45 ± 1 | 14.5 ± 1.31 |
| MMSE<sup>a,b,c</sup> | 29 ± 0.24 | 26.72 ± 0.46 | 21.86 ± 1.1 |
| ACE<sup>a,b,c</sup> | 94.78 ± 1 | 85.71 ± 1.61 | 66.46 ± 3.9 |
| MoCA<sup>a,b,c</sup> | 28.63 ± 0.38 | 24.72 ± 0.52 | 18.5 ± 1.33 |
| CDR global<sup>a,b,c</sup> | 0.21 ± 0.06 | 0.57 ± 0.05 | 1.17 ± 0.19 |

MMSE – Mini-mental state examination, ACE – Addenbrooke's cognitive examination, MoCA – Montreal cognitive assessment, CDR – clinical dementia rating
<sup>a</sup> Statistically significance (p<.05) between CN and AD dementia
<sup>b</sup> Statistically significance (p<.05) between CN and MCI
<sup>c</sup> Statistically significance (p<.05) between CN and MCI

### Experimental stimuli

Stimuli used in the task consisted of personally familiar names of places, events and people. To minimize the effects of memory disruptions on orientation testing, a set of personally familiar stimuli was obtained from each participant prior to testing. Participants were presented with a list of potential stimuli and for each were asked to approximate its location (for space stimuli) or year (for time stimuli). Failing to reference both the relevant region of the country and at least one nearby landmark (space) or misevaluating the correct year (time), resulted in the removal of the specific stimulus from further testing. In addition, participants were asked to generate a list of 8 close family members, 8 friends, and 8 acquaintances, which was corroborated with either a child or spouse (for additional details see supplementary materials).

### Experimental procedure - fMRI task

In the orientation task, participants were presented with pairs of familiar stimuli consisting of names of either two cities in Israel, two events, or two people, and were asked to determine which of the two is closer to them: geographically closer to their current location for cities, chronologically closer to the present time for events, or personally closer to them for people. To standardize experimental sessions based on personalized sets of stimuli, stimuli were split into three distance categories. Trials were generated by pairing stimuli from adjacent distance categories only. Stimuli were presented using the Presentation software (Version 18.0, Neurobehavioral Systems, Inc., Berkeley, CA; for additional details see supplementary materials).

Trials were presented in a randomized block design, with each block containing three consecutive trials belonging to a specific domain and distance category. Each trial was presented for a maximum of 10 seconds (5 TRs). Experiments consisted of four experimental runs, each containing 12 three-trial blocks in randomized order, balanced for both domain and distance categories. Additionally, participants performed a lexical control task in two additional separate runs. In the lexical control task, participants were presented with stimuli pairs from the same sets but were instructed to indicate which of the words contains the letter “A”. Stimuli were presented using the Presentation software (Version 18.0, Neurobehavioral Systems, Inc., Berkeley, CA). Prior to the experiment, A 5-minute training task containing different stimuli was administered. See supplementary materials for more details. The task was modeled on our previous studies of orientation (Dafni-Merom et al., 2019; Hayman & Arzy, 2021; Peer et al., 2015; Peters-Founshtein et al., 2018)

### Statistical analyses

Efficacy scores (ES) (Townsend & Ashby, 1983) were computed for each participant and domain separately by calculating the ratio between the success rate (SR) and mean response time (RT). A global ES was calculated for each participant by averaging the ESs across the three domains. Subsequently, mean ESs were compared across the three groups (AD dementia, MCI, CN) using analysis of variance (ANOVA) and Tukey-Kramer post-hoc tests. For the neuropsychological tests, scores were recorded according to the relevant testing guidelines.

### MRI and PET Data Acquisition and Preprocessing

For details regarding MRI and FDG-PET data acquisition and preprocessing please refer to supplementary materials.

### Voxel-based morphometry (VBM)

We applied voxel-based morphometry (VBM) analysis to compare gray matter (GM) density between CN, MCI and AD dementia groups (Ashburner & Friston, 2000). Specifically, we used a general linear model (GLM) (Worsley & Friston, 1995) to perform voxel-wise two-sample t-test for each of the clinical contrasts (CN–MCI, MCI–AD-dementia, CN–AD-dementia), with age, years of education, gender and total intracranial volume included as nuisance variables (see Supplementary Figure S1). Here, and in all further analyses we applied a false discovery rate (FDR) correction for multiple comparisons (P<0.05), and cluster size thresholding of 20 voxels. All analyses were performed using the statistical parametric mapping (SPM) 12 software package (version 7219), and in-house Matlab scripts (version 2019b, Mathworks, Natick, MA, USA). In-house scripts are publicly available (https://www.neuropsychiatrylab.com/codes).

### PET Analysis

To correct for AD-unrelated variance, FDG standardized uptake values (SUV) were first normalized by mean cerebellar GM SUV, to produce SUV ratio (SUVr) maps (Marcus, Mena, & Subramaniam, 2014). We then constructed a GLM to compare glucose uptake between the CN, MCI and AD dementia groups. Specifically, we performed voxel-wise two-sample t-tests comparing SUVr values between the three clinical contrasts (CN-MCI, MCI-AD-dementia, CN-AD-dementia), with age, years of education, and gender included as nuisance variables (Figure S1) (Kanda et al., 2008).

### Identification of orientation-evoked activity

To assess the selective activations elicited by different experimental conditions, we applied a group-level random-effects GLM analysis using data from all participants. To isolate orientation-specific activity, we contrasted the (1) spatial and temporal conditions (spatio-temporal) and, separately, the (2) social condition with the lexical control task (Peer et al., 2015).

### Orientation task analysis

We used a group-level random-effects GLM to compare spatio-temporal and social evoked activations between the CN, MCI, and AD dementia groups. Specifically, we performed voxel-wise two-sample t-tests comparing task parameter estimates of each domain in the three clinical contrasts with age, years of education, and gender included as nuisance variables. To exclude non-specific activations, maps of task over lexical control in all participants served as inclusive masks.

### Region-of-interest analysis

To associate task-evoked patterns of brain activity as well as glucose metabolism in the brain to DMN topology, ROIs for DMN subnetworks A, B, and C were extracted from the Schaefer 200 atlas (Schaefer et al., 2018). DMN ROIs were used to compare GM density, glucose uptake (Figure S2), and task evoked activity (parameter estimates of space-time and person over rest) between CN, MCI, and AD dementia groups. ANOVA and the Tukey-Kramer post-hoc test were used in all the comparisons.

### Mediation analysis

We set to test the hypothesis that neurodegeneration, expressed as changes in glucose metabolism, accounts for some of the shared variance between orientation-evoked activity and orientation performance along the AD continuum. For this purpose, we conducted a whole-brain voxel-wise mediation analysis using the Bootstrap Regression Analysis of Voxelwise Observations (BRAVO) toolbox (https://sites.google.com/site/bravotoolbox). Two three-path models were constructed, separately for spatio-temporal and social tasks (see Figure 4A), with parameter estimates for task-over-rest as the predictor variable (“X” in Figure 4A), task ES as outcome variable (“Y” in Figure 4A), and FDG-SUVr as mediator variable (“M” in Figure 4A). The mediation analysis tested whether the relation (path c) between the predictor variable (“X”, space-time/person beta) and an outcome variable (“Y”, space-time/person ES) is significantly attenuated when the relation between X and a mediator variable (“M”, FDG-SUVr) (path a) and the relation between M and Y (path b) are added to the model (Figure 4A). Mediation effect sizes were computed for every voxel. Significance was assessed through a permutation procedure with 10,000 iterations and corrected through FDR for voxel-wise multiple comparisons. Importantly, Models were applied to all participants and were agnostic to the clinical labels. To test the specificity of the mediation-related effects for the orientation tasks, we repeated the above analysis for the space-time and person domains of the lexical control task (Figure S3).

**Figure 1.**
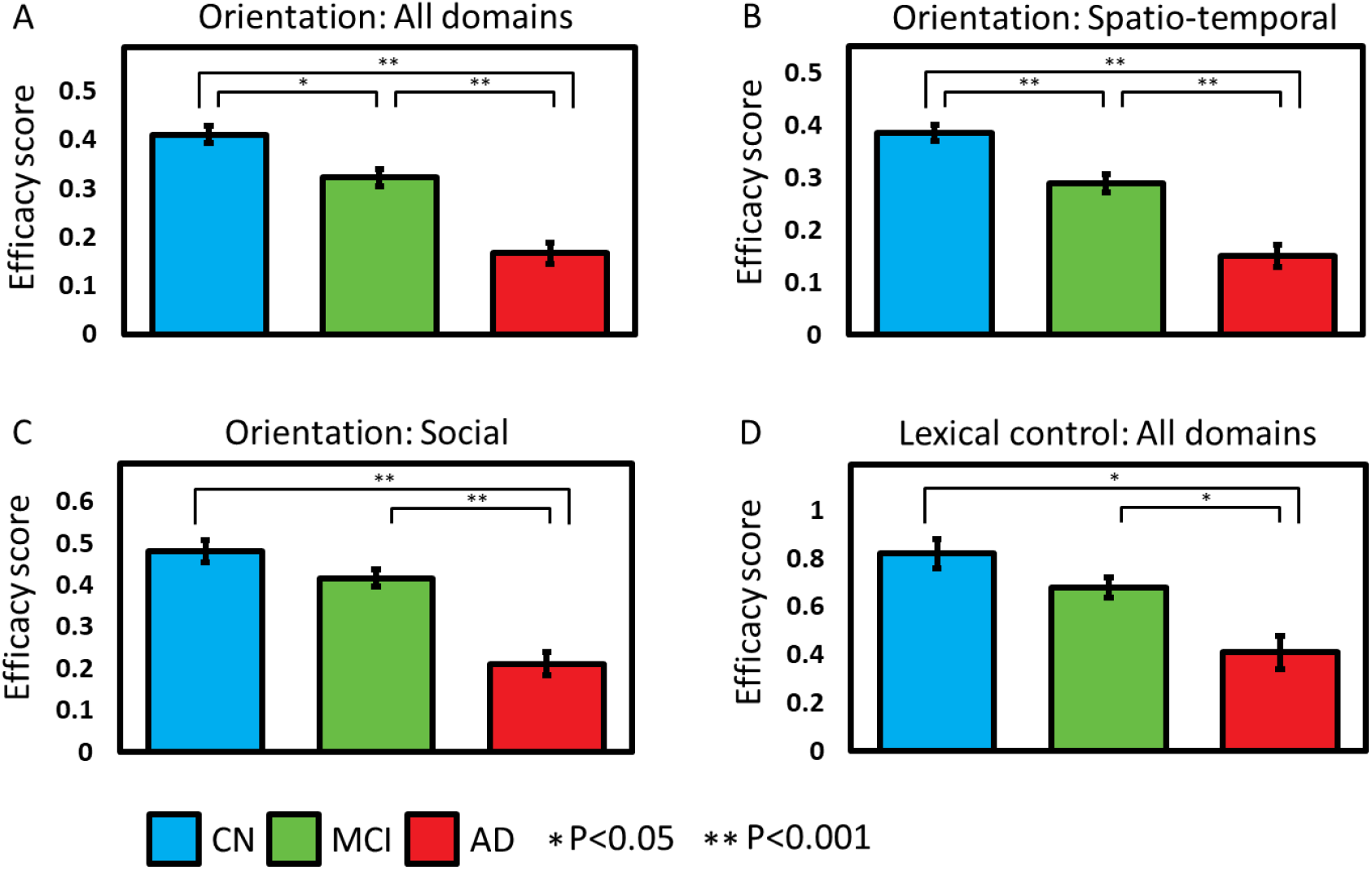
Spatio-temporal and social orientation changes along the AD continuum. Mean efficiency scores (ES) of CN (N= 16, blue), MCI (N = 23, green) and AD dementia participants (N = 12, red) for the orientation task in all domains (A), the orientation task in the domains of space and time (spatio-temporal) (B), the orientation task in person (social) (C), and the lexical control task in all domains (D). Significant CN-MCI differences were found in all domain orientation (A; P < 0.05), and space and time orientation (B; P < 0.001). Significant MCI-AD dementia and CN-AD dementia differences were found in all domains, spatio-temporal and social orientation task ES (A, B, C; P < 0.001), as well as in the lexical task (D; P < 0.05). Statistical significance was estimated using ANOVA and Tukey-Kramer post hoc test.

**Figure 2.**
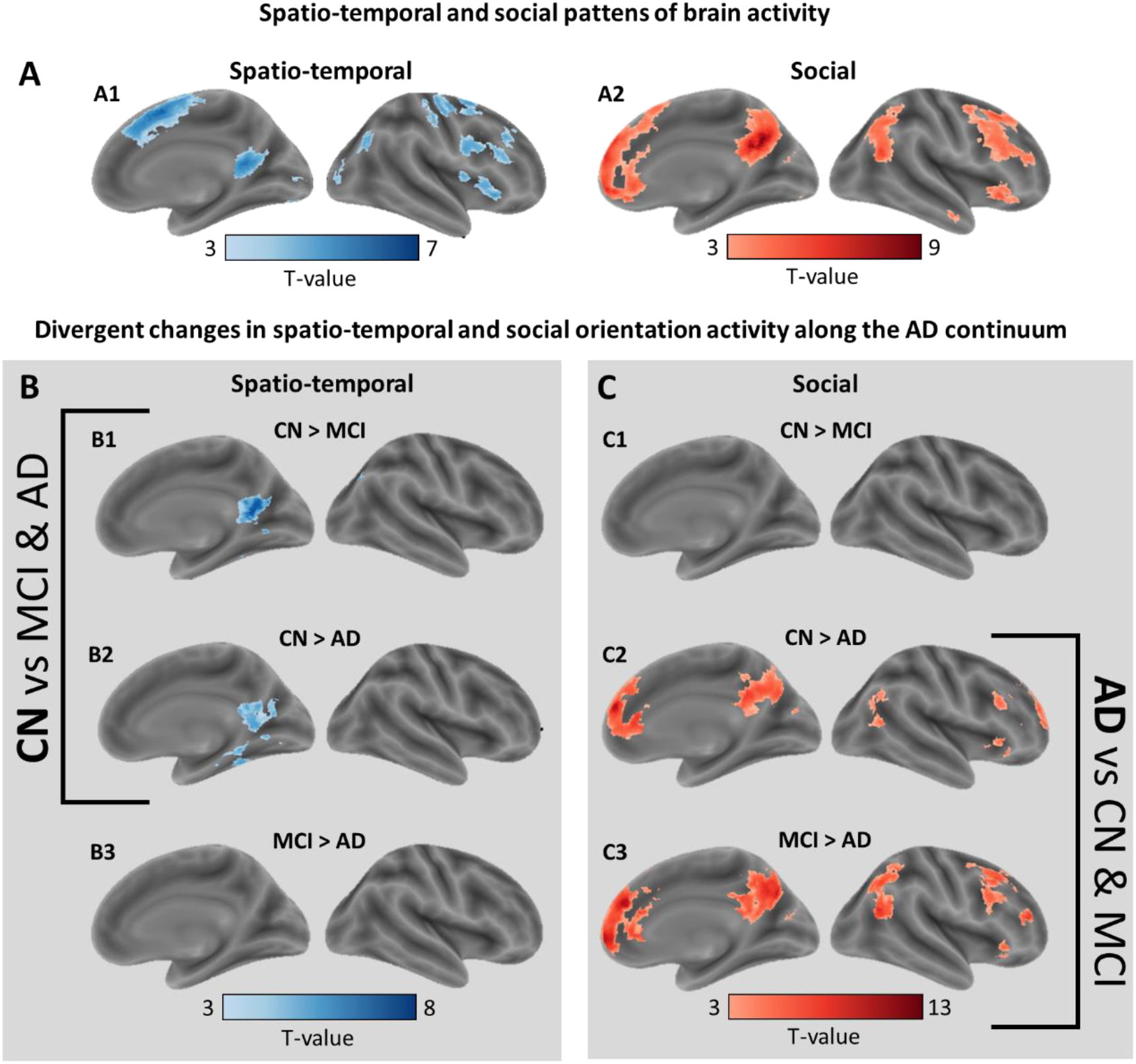
Changes in spatio-temporal and social orientation activity in AD. (A) Spatio-temporal and social pattens of brain activity. Results of GLM analysis exhibiting contrast maps of spatio-temporal (A1) and social (A2) orientation tasks over the lexical control task (All participants, DF=50, P<0.05 FDR corrected, cluster size > 20 voxels). (B, C) Divergent changes in spatio-temporal and social orientation activity. Spatio-temporal (B) and social (C) disorientation contrasted task-evoked activity maps across the three clinical groups: CN greater than MCI participants in spatio-temporal (B1) and social (C1) orientation (DF=38, P<0.05 FDR corrected, cluster size thresholding of 20 voxels); CN greater than AD dementia participants in spatio-temporal (B2) and social (C2) orientation (DF=34, P<0.05 FDR corrected, cluster size thresholding of 20 voxels); MCI greater than AD dementia participants in spatio-temporal (B3) and social (C3) orientation (DF=27, P<0.05 FDR corrected, cluster size thresholding of 20 voxels).

**Figure 3.**
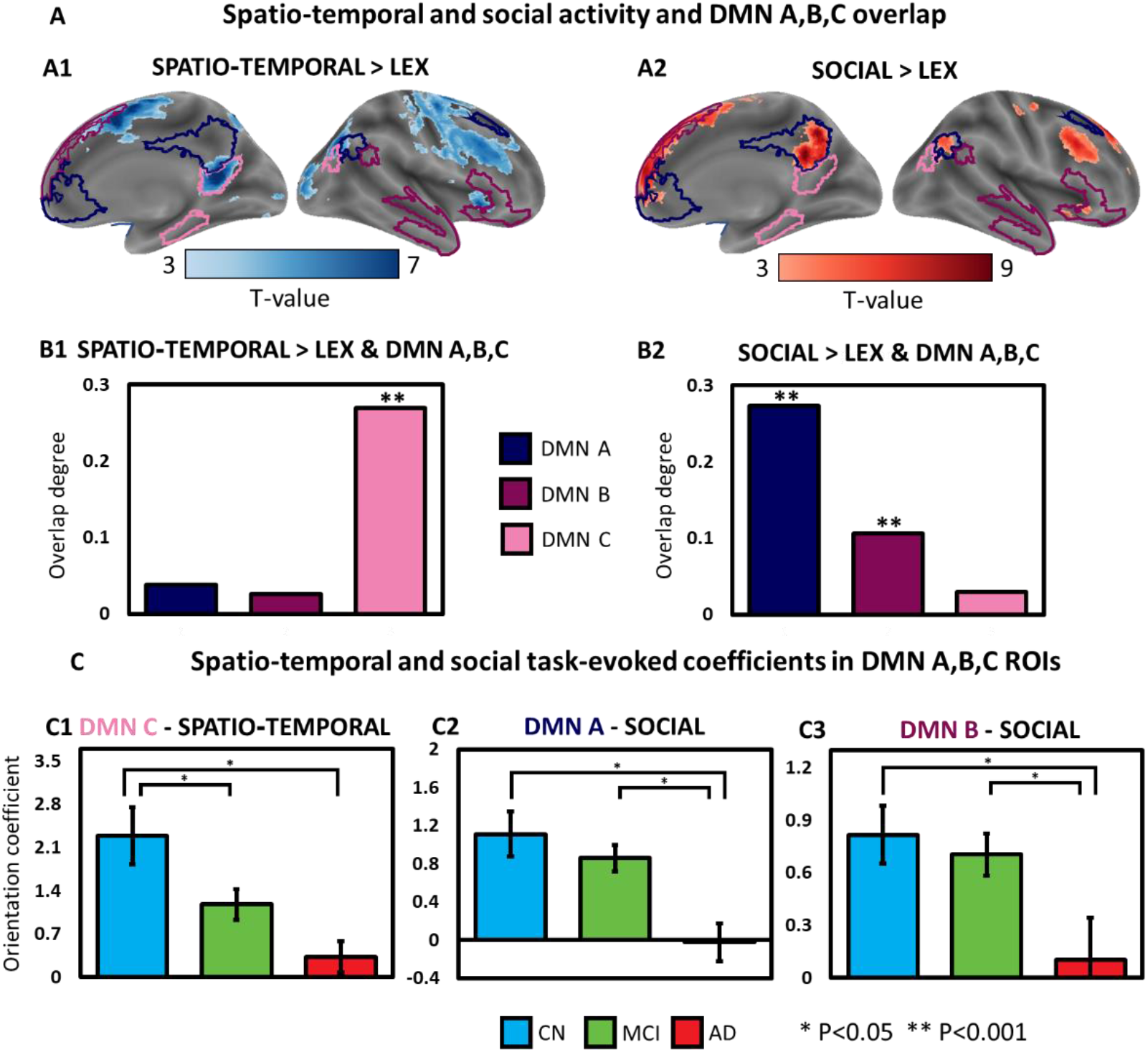
Orientation-evoked activity overlaps Default Mode sub-networks differently. **(A)** Spatio-temporal and social activity and DMN A, B, and C overlap. Delineations of DMN sub-networks (Schaefer et. al, 2018) DMN A (dark), DMN B (medium), DMN C (light) superimposed on maps of spatio-temporal (A1) and social (A2) orientation tasks (Orientation > Lexical control; All participants, DF=50, P<0.05 FDR corrected, cluster size > 20 voxels). **(B)** The precent of overlap between supra-threshold task-evoked spatio-temporal (B1) and social (B2) maps and DMN subnetworks A, B, and C. Asterisks indicate significant overlap (permutation test, 10,000 iterations). **(C)** Spatio-temporal and social task-evoked coefficients in DMN A, B, and C ROIs. Mean GLM-derived parameter estimates for social (C2 and 3) and spatio-temporal (C1) orientation (>rest) in significantly overlapping (B) DMN subnetworks (C1 – DMN C - spatio-temporal; C2–DMN B – social; C3 – DMN C - social) for CN (N= 16, blue), MCI (N = 23, green) and AD dementia (N = 12, red). Significant differences were found between CN and AD dementia participants in DMN A, B, and C, between MCI and AD dementia participants in DMN A and B, and between CN and MCI participants in DMN C (ANOVA and Tukey-Kramer post hoc test, P < 0.05).

**Figure 4.**
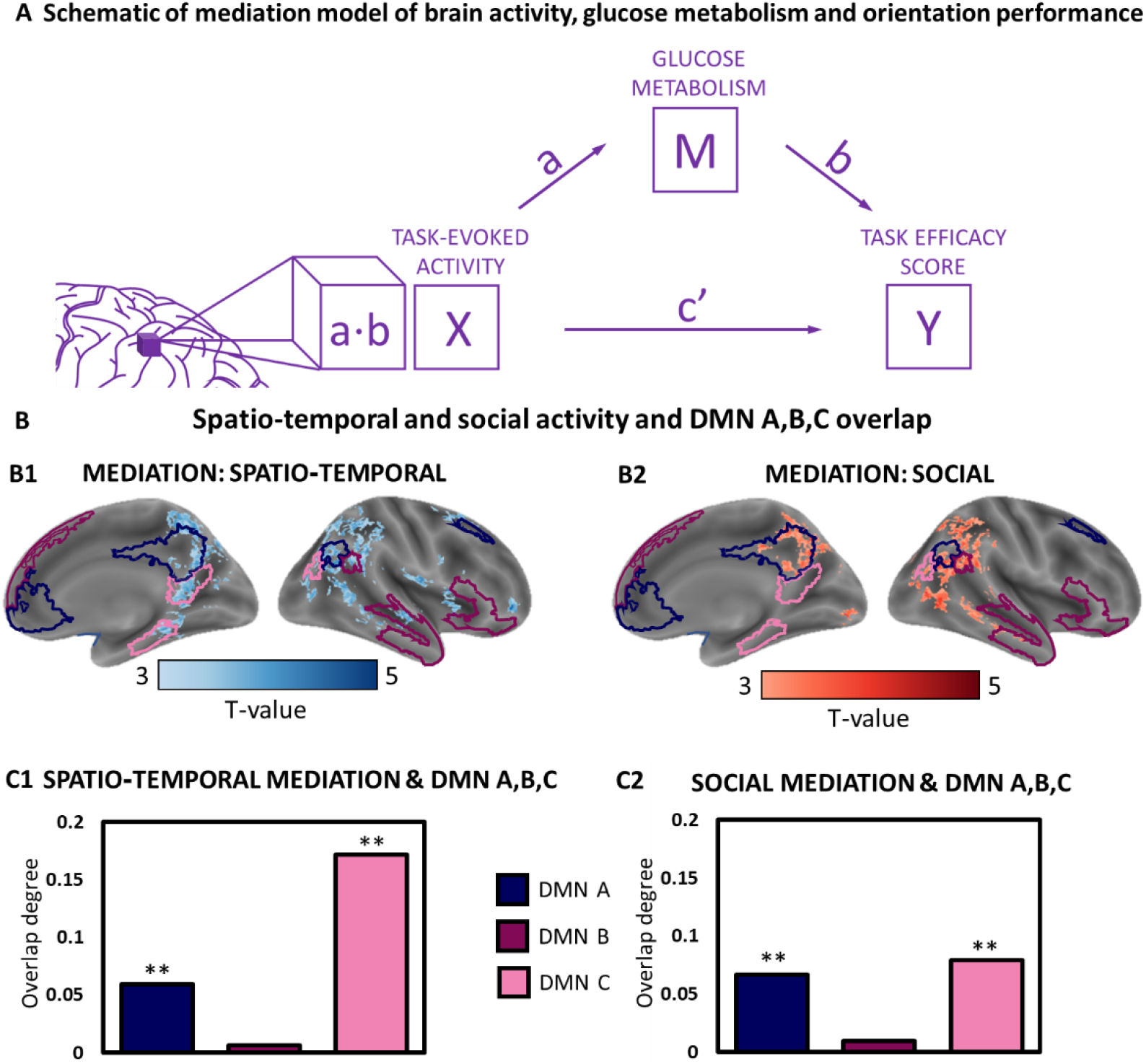
Mediation models of brain activity, glucose metabolism and orientation performance. **(A)** Mediation analysis was used to test the hypothesis that changes in FDG-PET uptake (M) across the AD continuum alter the relations between spatio-temporal (B2) and social (B2) orientation-evoked brain activity (X) and orientation task performance (Y). **(B)** Spatio-temporal and social mediation and DMN A, B, and C overlap. Delineations of DMN sub-networks (Schaefer et. al, 2018) DMN A (dark), DMN B (medium), and DMN C (light) superimposed on maps of spatio-temporal (B1) and social (B2) mediation (P<0.05, FDR-corrected). **(C)** The precent of overlap between supra-threshold task-evoked spatio-temporal (C1) and social (C2) suprathreshold mediation maps and DMN subnetworks A, B, and C. Asterisks indicate significant overlap (permutation test, 10,000 iterations).

### Permutation analysis for overlap significance

We performed overlap analysis to assess the correspondences between DMN-A, B, and C ROIs; spatio-temproal and social orientation-evoked maps of activity (Figure 3); spatio-temproal and social maps of mediation (Figure 4); and clinical contrasts for VBM (Figure S1) and FDG-SUVr (Figure S1). To assess overlap we quantified the shared number of suprathreshold (orientation, mediation, VBM, FDG) and DMN ROI voxels and divided by the total number of DMN ROI voxels. To determine significance of overlap we used a permutation analysis. We generated 10,000 permutation maps in which the same number of suprathreshold voxels was randomly shuffled, and then calculated the proportion of permutation maps in which the shared number of voxels was equal to or greater than the overlap between the original masks. This effectively determined the probability of observing the level of overlap we found with a random pattern of suprathreshold parameter.

## Results

### Spatio-temporal and social orientation performance is differently affected along the AD-continuum

Behavioral results for the orientation task across all domains showed significant differences between the 3 clinical groups (p<0.05, ANOVA and Tukey-Kramer post-hoc test). Participants with AD-dementia scored significantly lower than participants with MCI, and the latter—lower than CN participants (mean±SEM: 0.17±0.02[*sec^−1^*], 0.32±0.021[*sec^−1^*], 0.41±0.02[*sec^−1^*]; Figure 1A). Efficacy scores (ES) in the spatio-temporal domains (mean±SEM: 0.15±0.02[*sec^−1^*], 0.29±0.02[*sec^−1^*], 0.38±0.02[*sec^−1^*]; Figure 1B) showed significant differences between all 3 clinical groups (AD dementia, MCI, CN, respectively; P<0.05). ES in the social domain (mean±SEM: 0.21±0.03[*sec^−1^*], 0.42±0.02[*sec^−1^*], 0.48±0.02[*sec^−1^*]; Figure 1C) showed significant differences only between AD-dementia and non-AD participants (CN and MCI), comparable with the lexical control task (mean±SEM: 0.4±0.07[*sec^−1^*], 0.68±0.04[*sec^−1^*], 0.82±0.06 [*sec^−1^*]; Fig. 1D).

### Divergent changes in spatio-temporal and social activity along the AD continuum

Spatio-temporal orientation was shown to activate the precuneus, parieto-occipital sulcus, anterior and posterior cingulate cortices, parahippocampal and supramarginal gyri bilaterally, and the left superior frontal gyrus (Figure 2A). Social orientation activated the anterior and posterior cingulate cortices, and the angular and the superior medial gyri, and the putamen bilaterally (Figure 2A). Subsequent second level GLM analysis revealed significant differences in spatio-temporal orientation between CN and MCI participants and CN and AD-dementia participants in the precuneus, posterior cingulate cortex, parahippocampal gyri and hippocampus bilaterally (Figure 2B). Second level GLM analysis of social orientation showed significant differences between MCI and AD-dementia participants as well as CN and AD-dementia participants in the precuneus, superior medial and angular gyri bilaterally (Figure 2C).

### Spatio-temporal and social orientation-evoked activity overlap differently with Default Mode sub-networks

To examine whether discrete brain networks underlie spatio-temporal and social orientation we overlapped suprathreshold task-evoked activation maps with DMN A, B, and C ROIs. Spatio-temporal orientation activity was found to significantly overlap the DMN-C (27%, P<0.001, Figure 3A and B). Social orientation activations were found to significantly overlap the DMN-A (28%, P<0.001), and B (11%, P<0.001, 3A and B). We then compared mean task evoked coefficient estimated for spatio-temporal and social orientation among the CN, MCI and AD dementia groups, in each of the DMN ROIs. DMN ROI analysis for the spatio-temporal orientation showed differences between CN and MCI and AD-dementia participants in DMN-C only (P <0.05, Figure 3C and S2A). For social orientation, DMN ROI analysis showed differences between AD-dementia participants and MCI and CN participants in DMN A and B only (P<0.05, Figure 3C and S2B).

### AD-related hypometabolism mediates the relations between orientation performance and orientation-evoked activity

Spatio-temporal mediation effects were found to be significant (P<0.05, FDR-corrected) in the parahippocampal gyrus, posterior cingulate cortex and precuneus, significantly overlapping DMN-A (7%, P<0.001, Figure 4C) and DMN-C (17%, P<0.001, Figure 4C). Social mediation effects were found to be significant in the precuneus (P<0.05, FDR-corrected), significantly overlapping with DMN-A (7%, P<0.001, Figure 4C) and DMN-C (8%, P<0.001, Figure 4C). Mediation analysis for the lexical control task in space-time revealed small overlap with DMN-C (1%, P<0.001, Figure S3B and C). Social lexical mediation analysis revealed small overlap with DMN A (3%, P<0.001, Figure S3B2 and S3C2).

## Discussion

Our study revealed that an early stage of AD-related decline, MCI, manifests in spatio-temporal disorientation and task-evoked hypoactivation in temporoparietal regions, significantly overlapping the DMN-C subnetwork. Participants at the later stage of AD-dementia exhibited social disorientation and task-evoked hypoactivation in frontoparietal regions, significantly overlapping the DMN-A subnetwork. Moreover, the changes in task-evoked brain activity followed the pattern of glucose hypometabolism. Mediation analysis showed hypometabolism to mediate the relations between orientation-evoked activity and task performance along the AD continuum.

The DMN is a network of interconnected brain regions, which includes medial prefrontal, posterior cingulate and hippocampal brain structures. The DMN is known to activate when individuals engage in self-referential tasks, such as autobiographical memory retrieval and future planning (Buckner et al., 2008). In AD, early works have demonstrated a high degree of overlap between maps of DMN connectivity and patterns of structural and metabolic disruptions, as well as Aβ and tau deposition (Buckner et al., 2005; Hoenig et al., 2018; Palmqvist et al., 2017). More specifically, in AD patients, functionally connected regions were found to correlate with tau accumulation rates (Franzmeier, Neitzel, et al., 2020), corroborating the hypothesis that DMN connectivity facilitates trans-synaptic dispersion of tau across the brain (Adams et al., 2019; Buckner et al., 2005; Franzmeier, Dewenter, et al., 2020; Franzmeier, Neitzel, et al., 2020). Independently, several works have shown the DMN to comprise several distinct subnetworks (Andrews-Hanna et al., 2010; Buckner et al., 2008; Buckner & DiNicola, 2019). The detailed organization of these networks is revealed both in group and single subject level analyses. Evidence emerging from such studies suggests that the DMN comprises two or three separate networks with clear spatial distinctions along the posterior and anterior midline. Here we demonstrated the association between DMN subnetworks and spatio-temporal and social orientation in AD. Specifically, our findings suggest that disorientation, manifesting as progressive disruptions in behavioral performance and task-evoked brain activity is linked to sequential disruption in DMN-C (early) and DMN-A (late) regions along the AD continuum. Jointly, these results raise the possibility that DMN subnetworks are associated with distinct phases in AD progression.

Additional sources of evidence offer complementary accounts of the roles DMN subnetworks potentially play in AD pathology. PET neuroimaging of the two molecular AD hallmarks, Aβ and tau, has revealed distinct patterns of deposition across the brain. Aβ initially accumulates in medial parietal regions and spreads from neocortex to allocortex to brainstem. Tau, by contrast, first becomes evident in the entorhinal cortex, from which it spreads to limbic areas, and from there to the neocortex. In early stages of the disease, the pattern of Aβ deposition markedly overlaps with DMN A, while tau deposition markedly overlaps with DMN-C (van der Kant, Goldstein, & Ossenkoppele, 2020). Jointly, the associations between Aβ, tau and DMN subnetworks and our findings showing early DMN-C and late DMN-A hypometabolism, may suggest that at later stages of the disease, Aβ accumulation in DMN A facilitates the spread of tau beyond DMN C brain regions (Busche & Hyman, 2020). As Aβ was independently shown to affect functional connectivity (Lin et al., 2020), it is possible that Aβ felicitates tau spread beyond DMN-C regions by altering DMN A/C connectivity. This hypothesis is supported by the discovery of primary age-related tauopathy (PART), a neuropathological entity defined by the presence of tau in the absence of Aβ, which has been characterized by (1) tau remaining confined to the MTL regions and (2) appearing in cognitively intact older adults (Crary et al., 2014). As the nature of the relationship between Aβ and tau eludes consensus, the prospect of functional connectivity, specifically between subnetworks of the DMN, could potentially inform the mechanisms of propagation of AD neuropathology across the brain.

Our findings not only mark the significance of orientation testing in AD but also may suggest a role for AD as a potential neurodegenerative disease model of disorientation (Peer, Lyon, & Arzy, 2014). From this perspective, our results may contribute to the enduring question of domain-specific (B. Gauthier & van Wassenhove, 2016; Silson, Steel, Kidder, Gilmore, & Baker, 2019) versus domain-general (Park, Miller, & Boorman, 2021) systems of orientation. Coinciding with previously published findings (Kumaran & Maguire, 2005; Peer et al., 2015; Silson et al., 2019), here we linked spatio-temporal and social processing with temporoparietal, and frontoparietal regions, respectively. The diverging patterns of activity and vulnerability along the AD continuum for spatio-temporal and social orientation suggests a domain-dedicated framework, with its various components sequentially affected along the AD continuum.

However, several studies have challenged this paradigm of spatio-temporal and social dedicated brain regions, showing that, under some conditions, temporoparietal regions are involved in social-domain tasks, and frontoparietal regions - with spatial tasks. As a possible way of reconciling these seemingly contradictory findings, we propose that spatio-temporal and social activations represent special cases of archetypical modes of information processing (Kaplan & Friston, 2018; Peer et al., 2014; Whittington, McCaffary, Bakermans, & Behrens, 2022), underlied by temporoparietal (DMN-C) and frontoparietal (DMN-A) regions, respectively (Byrne, Becker, & Burgess, 2007; Whittington et al., 2020). Specifically, DMN-C regions were proposed to prioritize relational information, while DMN-A regions were found to accentuate self-referential aspects of the experience. In our task, relational and self-referential processes associated with spatio-temporal and social orientation, respectively, however under different task conditions these associations could potentially shift. In future studies we intend to generalize beyond specific tasks and define the underlying cognitive roles of the DMN subnetworks.

Social engagement in AD has been explored from several perspectives (Bennett, Schneider, Tang, Arnold, & Wilson, 2006; Fredericks et al., 2018; Sabat & Gladstone, 2010; Sabat & Lee, 2012; Sturm et al., 2013; Wilson et al., 2007). One approach focuses on disruptions in social mapping and orientation, i.e the ability to mentalize relational representation of other people within a multi-dimensional (social) space (Schafer & Schiller, 2018). Here we demonstrate the relative resilience of social orientation, which finally breaks down in the later stages of AD, and its association to the DMN-A subnetwork. Brain systems, such as hippocampal place cells or entorhinal grid cells, once considered dedicated to spatial computations, are gradually recognized for their role in social cognition (Omer, Maimon, Las, & Ulanovsky, 2018; Park et al., 2021). In an aforementioned study, Tavares and colleagues have shown that hippocampal BOLD signal during social “navigation” in the dimensions of power and affiliation was negatively correlated with social avoidance and neuroticism, suggesting a link between social orientation and interpersonal traits. Here we present evidence suggesting that a relative preservation of social orientation in early stages of AD (followed by late disruption) is linked to relatively late neurodegeneration in the DMN-A subnetwork regions. In the context of studies mapping social orientation to MTL structures, our results suggest that in addition to previously reported MTL regions, frontoparietal DMN-A regions play a critical role in social orientation.

To the best of our knowledge, ours is the first study to utilize a combination of PET and MR imaging to simultaneously assess neurodegeneration, task-induced brain activity, and performance within a single cohort. We used mediation analysis to jointly analyze functional, metabolic, and behavioral data, to examine whether (and where in the brain) hypometabolism mediates the relationship between orientation-evoked activity and task performance along the AD continuum. We conducted mediation analyses separately for spatio-temporal and social orientation, as well as for a lexical control task, matching the orientation task in format and stimuli yet differing in its cognitive requirements. For spatio-temporal and social orientation, mediation effects were found to be significant in DMN C and DMN A medial regions, respectively, suggesting AD-related neurodegeneration induces changes beyond simply suppressing activity and disrupting performance, but rather affecting the relations between activity and performance (Huijbers et al., 2015). Additional studies have shown mixed patterns of hypo- and hyperactivation in patients along the AD spectrum (Foster, Kennedy, Horn, Hoagey, & Rodrigue, 2018; R. Sperling, 2007). In one such study, Kunz and colleagues (Kunz et al., 2015) demonstrated that young asymptomatic APOE-ε4 carriers (AD risk multiplier) exhibit decreased entorhinal and increased hippocampal activity while navigating a 3D arena. Exploring later, clinically detectable, stages of AD, O’Brien and colleagues (O’Brien et al., 2010) administered a memory-encoding task to a group of MCI and older adult control participants while undergoing fMRI. The results demonstrated that while both groups were similarly successful in recall, stronger hippocampal activation in the MCI group during encoding correlated to a higher rate of cognitive decline, and sequential hypoactivation in follow-up scans. The effects of AD on brain activity therefore appear to relate both to the stage of the disease and to the task itself. In future efforts, we intend to model relationships between brain activity and behavior with specific neuroimaging markers of AD-pathology.

Our results and conclusion notwithstanding, this work is not free from limitations. First, in recent years there has been a push for progressively shifting the definition of AD from a syndromal to a biological one (Jack et al., 2018). Specifically, AD biomarkers, including Aβ, tau and neurodegeneration have been reorganized into the ATN framework (Kern et al., 2018). In the current study we focused on 2 separate markers of neurodegeneration: structural MRI and FDG-PET. In future studies we intend to incorporate Aβ and tau PET imaging in order to shift the focus from networks and activity to pathology. Additionally, the current study was cross-sectional and had a relatively small sample size, especially when considering the inclusion of 3 diagnostic groups (CN, MCI, and AD dementia). Future studies will consist of larger sample sizes and longitudinal follow-up, allowing us to better understand the directionality of these brain-behavior associations and their evolution over time in AD.

In conclusion, this study demonstrates the central role of disorientation in AD, and specifically the potential of evaluating orientation in multiple domains to differentiate between disease stages. We suggest that the relative early vulnerability of spatial and temporal orientation (compared to social orientation) stems from its association with temporoparietal regions, which are affected in early stages of the disease. Comparably, the relative resilience of social orientation stems from its association with the frontoparietal cortex, which is affected only in later stages of the disease. We attribute these associations to distinct underlying computations performed by functionally distinct subnetworks of the DMN. We suggest that in parallel with the rapidly evolving evidence in support of usage of biomarkers in AD diagnosis, establishing a data-driven neurocognitive profile of AD degeneration will greatly benefit disease diagnosis, monitoring and evaluation of treatment response.

## Acknowledgements

We are grateful to Zohar Nitsan, Inbal Machcat and Hagai Baruch for their help in participant scanning, and to Dr. Limor Shaharabani-Gargir for her invaluable help with participant management. To Dr. Michael Peer, Dr. Moshe Roseman and Mr. Amnon Dafni-Merom for their insightful comments. This work was supported by the Israeli Science foundation (grant no. 3213/19) and the NIH (grant no. R21 AG070877 to GAM, HS and SA). SA is grateful to Mr. Ronald Abramson and The Glazer foundation for their generous support.

## References

Adams, J. N., Maass, A., Harrison, T. M., Baker, S. L., & Jagust, W. J. (2019). Cortical tau deposition follows patterns of entorhinal functional connectivity in aging. ELife, 8, 1–22. https://doi.org/10.7554/eLife.49132

Albert, M. S., DeKosky, S. T., Dickson, D., Dubois, B., Feldman, H. H., Fox, N. C.,… Phelps, C. H. (2011). The diagnosis of mild cognitive impairment due to Alzheimer’s disease: Recommendations from the National Institute on Aging-Alzheimer’s Association workgroups on diagnostic guidelines for Alzheimer’s disease. Alzheimer’s and Dementia, 7(3), 270–279. https://doi.org/10.1016/j.jalz.2011.03.008

Andrews-Hanna, J. R., Reidler, J. S., Sepulcre, J., Poulin, R., & Buckner, R. L. (2010). Functional-Anatomic Fractionation of the Brain’s Default Network. Neuron, 65(4), 550–562. https://doi.org/10.1016/j.neuron.2010.02.005

Ashburner, J., & Friston, K. J. (2000). Voxel-based morphometry - The methods. NeuroImage, 11(6 I), 805–821. https://doi.org/10.1006/nimg.2000.0582

Barnett, A. J., Reilly, W., Dimsdale-Zucker, H., Mizrak, E., Reagh, Z., & Ranganath, C. (2020). Organization of cortico-hippocampal networks in the human brain. In bioRxiv. https://doi.org/10.1101/2020.06.09.142166

Barnett, A. J., Reilly, W., Dimsdale-Zucker, H. R., Mizrak, E., Reagh, Z., & Ranganath, C. (2021). Intrinsic connectivity reveals functionally distinct cortico-hippocampal networks in the human brain. In PLoS Biology (Vol. 19). https://doi.org/10.1371/journal.pbio.3001275

Barnett, D. A., Arnold, S. E., Valenzuela, M. J., Brayne, C., & Schneider, J. A. (2014). Cognition in Late Life. Acta Neuropathol., 127(1), 137–150. https://doi.org/10.1007/s00401-013-1226-2.Cognitive

Bennett, D. A., Schneider, J. A., Tang, Y., Arnold, S. E., & Wilson, R. S. (2006). The effect of social networks on the relation between Alzheimer’s disease pathology and level of cognitive function in old people: a longitudinal cohort study. Lancet Neurology, 5(5), 406–412. https://doi.org/10.1016/S1474-4422(06)70417-3

Berrios, G. E. (1982). Disorientation states and psychiatry. Comprehensive Psychiatry, 23(5), 479–491. https://doi.org/10.1016/0010-440X(82)90161-4

Buckner, R. L., Andrews-Hanna, J. R., & Schacter, D. L. (2008). The brain’s default network: Anatomy, function, and relevance to disease. Annals of the New York Academy of Sciences, 1124, 1–38. https://doi.org/10.1196/annals.1440.011

Buckner, R. L., & DiNicola, L. M. (2019). The brain’s default network: updated anatomy, physiology and evolving insights. Nature Reviews Neuroscience, 20(10), 593–608. https://doi.org/10.1038/s41583-019-0212-7

Buckner, R. L., Snyder, A. Z., Shannon, B. J., Larossa, G., Sachs, R., Fotenos, A. F.,… Mintun, M. A. (2005). Molecular, Structural, and Functional Characterization of Alzheimer’s Disease: Evidence for a Relationship between Default Activity, Amyloid, and Memory. Neuroscience, 25(34), 7709–7717. https://doi.org/10.1523/JNEUROSCI.2177-05.2005

Busche, M. A., & Hyman, B. T. (2020). Synergy between amyloid-β and tau in Alzheimer’s disease. Nature Neuroscience, 23(10), 1183–1193. https://doi.org/10.1038/s41593-020-0687-6

Byrne, P., Becker, S., & Burgess, N. (2007). Remembering the past and imagining the future: a neural model of spatial memory and imagery. Psychological Review, 114(2), 340–375. https://doi.org/10.1037/0033-295X.114.2.340

Calhoun, V. D., & Sui, J. (2016). Multimodal Fusion of Brain Imaging Data: A Key to Finding the Missing Link(s) in Complex Mental Illness. Biological Psychiatry: Cognitive Neuroscience and Neuroimaging, 1(3), 230–244. https://doi.org/10.1016/j.bpsc.2015.12.005

Coughlan, G., Coutrot, A., Khondoker, M., Minihane, A. M., Spiers, H., & Hornberger, M. (2019). Toward personalized cognitive diagnostics of at-genetic-risk Alzheimer’s disease. Proceedings of the National Academy of Sciences of the United States of America, 116(19), 9285–9292. https://doi.org/10.1073/pnas.1901600116

Coughlan, G., Laczó, J., Hort, J., Minihane, A. M., & Hornberger, M. (2018, August 6). Spatial navigation deficits — Overlooked cognitive marker for preclinical Alzheimer disease? Nature Reviews Neurology, Vol. 14, pp. 496–506. https://doi.org/10.1038/s41582-018-0031-x

Crary, J. F., Trojanowski, J. Q., Schneider, J. A., Abisambra, J. F., Abner, E. L., Alafuzoff, I.,… Nelson, P. T. (2014). Primary age-related tauopathy (PART): a common pathology associated with human aging. Acta Neuropathologica, 128(6), 755–766. https://doi.org/10.1007/s00401-014-1349-0

Dafni-Merom, A., Peters-Founshtein, G., Kahana-Merhavi, S., & Arzy, S. (2019). A unified brain system of orientation and its disruption in Alzheimer’s disease. Annals of Clinical and Translational Neurology, 6(12), 2468–2478. https://doi.org/10.1002/acn3.50940

DeIpolyi, A. R., Rankin, K. P., Mucke, L., Miller, B. L., Gorno-Tempini, M. L. (2007). Spatial cognition and the human navigationnetwork in AD and MCI. Neurology, 69, 986–997.

Du, M., Basyouni, R., & Parkinson, C. (2021). How does the brain navigate knowledge of social relations? Testing for shared neural mechanisms for shifting attention in space and social knowledge. NeuroImage, 235(March), 118019. https://doi.org/10.1016/j.neuroimage.2021.118019

Dubois, B., Slachevsky, a, Litvan, I., & Pillon, B. (2000). The FAB: a Frontal Assessment Battery at bedside. Neurology, 55(11), 1621–1626. https://doi.org/10.1212/WNL.57.3.565

El Haj, M., & Antoine, P. (2018). Context Memory in Alzheimer’s Disease: The “who, Where, and When.” Archives of Clinical Neuropsychology, 33(2), 158–167. https://doi.org/10.1093/arclin/acx062

Franzmeier, N., Dewenter, A., Frontzkowski, L., Dichgans, M., Rubinski, A., Neitzel, J.,… Ewers, M. (2020). Patient-centered connectivity-based prediction of tau pathology spread in Alzheimer’s disease. (November).

Franzmeier, N., Neitzel, J., Rubinski, A., Smith, R., Strandberg, O., Ossenkoppele, R.,… Raj, B. A. (2020). Functional brain architecture is associated with the rate of tau accumulation in Alzheimer’s disease. Nature Communications, 11(1), 1–17. https://doi.org/10.1038/s41467-019-14159-1

Fredericks, C. A., Sturm, V. E., Brown, J. A., Hua, A. Y., Bilgel, M., Wong, D. F.,… Seeley, W. W. (2018). Early affective changes and increased connectivity in preclinical Alzheimer’s disease. Alzheimer’s and Dementia: Diagnosis, Assessment and Disease Monitoring, 10, 471–479. https://doi.org/10.1016/j.dadm.2018.06.002

Gauthier, B., & van Wassenhove, V. (2016). Time is not space: Core computations and domain-specific networks for mental travels. Journal of Neuroscience, 36(47), 11891–11903. https://doi.org/10.1523/JNEUROSCI.1400-16.2016

Gauthier, S., Reisberg, B., Zaudig, M., Petersen, R. C., Ritchie, K., Broich, K.,… Winblad, B. (2006). Mild cognitive impairment. Lancet (London, England), 367(9518), 1262–1270. https://doi.org/10.1016/S0140-6736(06)68542-5

Hachinski, V. C., Iliff, L. D., Zilhka, E., Du Boulay, G. H., McAllister, V. L., Marshall, J.,… Symon, L. (1975). Cerebral blood flow in dementia. Archives of Neurology, 32(9), 632–637.

Hayman, M., & Arzy, S. (2021). Mental travel in the person domain. Journal of Neurophysiology, 20(7), 464–476. https://doi.org/10.1152/jn.00695.2020

Hoenig, M. C., Bischof, G. N., Seemiller, J., Hammes, J., Kukolja, J., Onur, Ö. A.,… Drzezga, A. (2018). Networks of tau distribution in Alzheimer’s disease. Brain, 141(2), 568–581. https://doi.org/10.1093/brain/awx353

Huijbers, W., Mormino, E. C., Schultz, A. P., Wigman, S., Ward, A. M., Larvie, M.,… Sperling, R. A. (2015). Amyloid-β deposition in mild cognitive impairment is associated with increased hippocampal activity, atrophy and clinical progression. Brain, 138(4), 1023–1035. https://doi.org/10.1093/brain/awv007

Kanda, T., Ishii, K., Uemura, T., Miyamoto, N., Yoshikawa, T., Kono, A. K., & Mori, E. (2008). Comparison of grey matter and metabolic reductions in frontotemporal dementia using FDG-PET and voxel-based morphometric MR studies. European Journal of Nuclear Medicine and Molecular Imaging, 35(12), 2227–2234. https://doi.org/10.1007/s00259-008-0871-5

Kaplan, R., & Friston, K. J. (2018). Hippocampal-entorhinal transformations in abstract frames of reference. Entorhinal Transformations in Abstract Frames of Reference, 414524. https://doi.org/10.1101/414524

Kern, S., Zetterberg, H., Kern, J., Zettergren, A., Waern, M., Höglund, K.,… Skoog, I. (2018). Prevalence of preclinical Alzheimer disease: Comparison of current classification systems. Neurology, 90(19), E1682–E1691. https://doi.org/10.1212/WNL.0000000000005476

Kumaran, D., & Maguire, E. A. (2005). The human hippocampus: Cognitive maps or relational memory? Journal of Neuroscience, 25(31), 7254–7259. https://doi.org/10.1523/JNEUROSCI.1103-05.2005

Kunz, L., Schröder, T. N., Lee, H., Montag, C., Lachmann, B., Sariyska, R.,… Axmacher, N. (2015). Reduced grid-cell-like representations in adults at genetic risk for Alzheimer’s disease. Science (New York, N.Y.), 350(6259), 430–433. https://doi.org/10.1126/science.aac8128

Lin, C., Ly, M., Karim, H. T., Wei, W., Snitz, B. E., Klunk, W. E., & Aizenstein, H. J. (2020). The effect of amyloid deposition on longitudinal resting-state functional connectivity in cognitively normal older adults. Alzheimer’s Research and Therapy, 12(1), 1–10. https://doi.org/10.1186/s13195-019-0573-1

Mathuranath, P. S., Nestor, P. J., Berrios, G. E., Rakowicz, W., & Hodges, J. R. (2000). A brief cognitive test battery to differentiate Alzheimer’s disease and frontotemporal dementia. Neurology, 55(11), 1613–1620. Retrieved from http://www.ncbi.nlm.nih.gov/pubmed/11113213

Mckhann, G., Drachman, D., Folstein, M., Katzman, R., Price, D., & Stadlan, E. M. (n.d.). Clinical diagnosis of Alzheimer’s disease: Task Force on Alzheimer’s Disease.

Morris, J. C. (1993). The Clinical Dementia Rating (CDR): current version and scoring rules. Neurology, 43(11), 2412–2414. https://doi.org/10.1212/WNL.43.11.2412-a

Nasreddine, Z., Phillips, N., Bédirian, V., Charbonneau, S., Whitehead, V., Colllin, I.,… Chertkow, H. (2005). The Montreal Cognitive Assessment, MoCA: a brief screening tool for mild cognitive impairment. Journal of the American Geriatrics Society, 53(4), 695–699. https://doi.org/10.1111/j.1532-5415.2005.53221.x

Omer, D. B., Maimon, S. R., Las, L., & Ulanovsky, N. (2018). Social place-cells in the bat hippocampus. Science, 359(6372), 218–224. https://doi.org/10.1126/science.aao3474

Palmqvist, S., Schöll, M., Strandberg, O., Mattsson, N., Stomrud, E., Zetterberg, H.,… Hansson, O. (2017). Earliest accumulation of β-amyloid occurs within the default-mode network and concurrently affects brain connectivity. Nature Communications, 8(1). https://doi.org/10.1038/s41467-017-01150-x

Park, S. A., Miller, D. S., & Boorman, E. D. (2021). Inferences on a multidimensional social hierarchy use a grid-like code. Nature Neuroscience, 24(9), 1292–1301. https://doi.org/10.1038/s41593-021-00916-3

Parkinson, C., Liu, S., & Wheatley, T. (2014). A common cortical metric for spatial, temporal, and social distance. The Journal of Neuroscience, 34(5), 1979–1987. https://doi.org/10.1523/JNEUROSCI.2159-13.2014

Peer, M., Lyon, R., & Arzy, S. (2014). Orientation and disorientation: Lessons from patients with epilepsy. Epilepsy and Behavior, 41, 149–157. https://doi.org/10.1016/j.yebeh.2014.09.055

Peer, M., Salomon, R., Goldberg, I., Blanke, O., & Arzy, S. (2015). Brain system for mental orientation in space, time, and person. Proceedings of the National Academy of Sciences, 112(35), 11072–11077. https://doi.org/10.1073/pnas.1504242112

Peters-Founshtein, G., Peer, M., Rein, Y., Kahana Merhavi, S., Meiner, Z., & Arzy, S. (2018). Mental-orientation: A new approach to assessing patients across the Alzheimer’s disease spectrum. Neuropsychology. https://doi.org/10.1037/neu0000463

Peters-founshtein, G., Peer, M., Rein, Y., Merhavi, S. K., Meiner, Z., Peters-founshtein, G.,… Meiner, Z. (2018). Mental-orientation: A new approach to assessing patients across the Alzheimer’s disease spectrum. Neuropsychology.

Petersen, R. C., Smith, G. E., Waring, S. C., Ivnik, R. J., Tangalos, E. G., & Kokmen, E. (1999). Mild Cognitive Impairment. Archives of Neurology, 56(3), 303. https://doi.org/10.1001/archneur.56.3.303

Sabat, S. R., & Gladstone, C. M. (2010). What intact social cognition and social behavior reveal about cognition in the moderate stage of Alzheimer’s disease: A case study. Dementia, 9(1), 61–78. https://doi.org/10.1177/1471301210364450

Sabat, S. R., & Lee, J. M. (2012). Relatedness among people diagnosed with dementia: Social cognition and the possibility of friendship. Dementia, 11(3), 315–327. https://doi.org/10.1177/1471301211421069

Schaefer, A., Kong, R., Gordon, E. M., Laumann, T. O., Zuo, X.-N., Holmes, A. J.,… Yeo, B. T. T. (2018). Local-Global Parcellation of the Human Cerebral Cortex from Intrinsic Functional Connectivity MRI. Cerebral Cortex, 28(9), 3095–3114. https://doi.org/10.1093/cercor/bhx179

Schafer, M., & Schiller, D. (2018). Navigating Social Space. Neuron, 100(2), 476–489. https://doi.org/10.1016/j.neuron.2018.10.006

Silson, E. H., Steel, A., Kidder, A., Gilmore, A. W., & Baker, C. I. (2019). Distinct subdivisions of human medial parietal cortex support recollection of people and places. ELife, 1–25. https://doi.org/10.1101/554915

Sturm, V. E., Yokoyama, J. S., Seeley, W. W., Kramer, J. H., Miller, B. L., & Rankin, K. P. (2013). Heightened emotional contagion in mild cognitive impairment and Alzheimer’s disease is associated with temporal lobe degeneration. Proceedings of the National Academy of Sciences of the United States of America, 110(24), 9944–9949. https://doi.org/10.1073/pnas.1301119110

Townsend, J. T., & Ashby, F. G. (1983). Stochastic Modeling of Elementary Psychological Processes. The American Journal of Psychology, 480. https://doi.org/10.2307/1422636

van der Kant, R., Goldstein, L. S. B., & Ossenkoppele, R. (2020). Amyloid-β-independent regulators of tau pathology in Alzheimer disease. Nature Reviews Neuroscience, Vol. 21, pp. 21–35. https://doi.org/10.1038/s41583-019-0240-3

Whittington, J. C. R., McCaffary, D., Bakermans, J. J. W., & Behrens, T. E. J. (2022). How to build a cognitive map. Nature Neuroscience, 25(10), 1257–1272. https://doi.org/10.1038/s41593-022-01153-y

Whittington, J. C. R., Muller, T. H., Mark, S., Chen, G., Barry, C., Burgess, N., & Behrens, T. E. J. (2020). The Tolman-Eichenbaum Machine: Unifying Space and Relational Memory through Generalization in the Hippocampal Formation. Cell, 183(5), 1249–1263.e23. https://doi.org/10.1016/j.cell.2020.10.024

Wilson, R. S., Krueger, K. R., Arnold, S. E., Schneider, J. A., Kelly, J. F., Barnes, L. L.,… Bennett, D. A. (2007). Loneliness and risk of Alzheimer disease. Archives of General Psychiatry, 64(2), 234–240. https://doi.org/10.1001/archpsyc.64.2.234

Worsley, K. J., & Friston, K. J. (1995). Analysis of fMRI time-series revisited — Again. NeuroImage, 2(3), 173–181. https://doi.org/10.1006/nimg.1995.1023

