## Supplementary Materials for "Lost in space(s): multimodal neuroimaging of disorientation along the Alzheimer’s disease continuum"

#### Supplementary Methods

*Obtaining Stimuli*

To obtain participant-tailored stimuli, prior to preforming the task, participants were presented with a list of potential stimuli. Space stimuli consisted of names of cities in Israel, distanced 8–150 km from participants' location. For each space stimulus, participants were asked to approximate its location (general region, relation to landmarks). Failing to reference both the relevant region of the country and at least one nearby landmark (space) resulted in the removal of the specific stimulus from further testing. Time stimuli consisted of two-word descriptions of common past events from personal life (e.g., first grandchild) or non-personal world events (e.g., Obama's inauguration). For each time stimulus, participants were asked to approximate the year it occurred. Misevaluating by more than five years (time), resulted in the removal of the specific stimulus from further testing. Person stimuli consisted of people's full names; participants were requested to generate a list of 8 close family members, 8 friends, and 8 acquaintances. Stimuli in each domain were assigned to one of three distance categories relative to the participants' own self-location. Stimuli of a personal nature were corroborated by an informant (family member—child or spouse).

*MRI Data Acquisition*

All participants were scanned using a Siemens® Biograph PET-MRI 3T scanner system with a 32-channel head coil (Siemens Medical Solutions, Erlangen, Germany). Blood oxygen level dependent (BOLD) fMRI was acquired using a whole-brain, gradient-echo (GE) echoplanar (EPI) [repetition time (TR)/echo time (TE) = 2,020/30 ms, flip angle = 75, field of view (FOV) = 192 × 192 mm, matrix = 64 × 64, 36 axial slices, slice thickness/gap = 4 mm/0.8 mm. In addition, high resolution (1 × 1 × 1 mm) T1-weighted anatomical images were acquired to aid spatial normalization to standard atlas space. The anatomic reference volume was acquired along the same orientation as the functional images [TR/TE = 2,000/2.34 ms, matrix = 256 × 256, 160 axial slices, 1-mm slice thickness, inversion time (TI) = 900 ms].

*MRI Preprocessing*

fMRI data were analyzed using the statistical parametric mapping (SPM) 12 software package, version 7487 and in-house Matlab (Mathworks) scripts (available at neuropsychiatrylab.com). Preprocessing of functional scans included slice-time correction (sinc interpolation), 3D motion correction by realignment to the first run image (2nd degree B-Spline interpolation), smoothing (full width at half maximum (FWHM) = 4 mm), and co-registration to the anatomical T1 images. Anatomical brain images were corrected for signal inhomogeneity, skull-stripped, and segmented into three different tissue compartments (gray matter, white matter, and cerebrospinal fluid (CSF)). All images were subsequently normalized to Montreal Neurological Institute (MNI) space (3 × 3 × 3 mm functional resolution, 4th degree B-Spline interpolation).

*PET-FDG Data Acquisition*

All participants were scanned using a Siemens® Biograph PET-MRI 3T scanner. The PET data were acquired in 3-dimensional mode, yielding a 155-mm field of view. For each [$^{18}$F]-Fluorodeoxyglucose (FDG)-PET acquisition, 17 frames (10 x 30 seconds and 7 x 300 seconds) were acquired over 45 minutes. A mean FDG sum image was created and used for subsequent analysis. Participants fasted for 4 hours preceding the FDG-PET scan. The mean injected dose for each tracer was 196.82 ± 7.6 MBq.

*FDG-PET Preprocessing*

FDG-PET standardized uptake value (SUV) images were coregistered to the participant's T1 image in native space, and subsequently normalized to MNI space, applying the parameters derived from the T1 normalization process. Normalized FDG-PET images were standardized, by dividing SUV values in each voxel by the mean cerebellar grey matter SUV, to produce SUV ratio (SUVr) maps. All images were then smoothed using an 8-mm Gaussian smoothing kernel.

*fMRI task procedure*

In the orientation task, participants were presented with pairs of familiar stimuli consisting of names of either two cities in Israel, two events, or two people, and were asked to determine which of the two is closer to them: geographically closer to their current location for cities, chronologically closer to the present time for events, or personally closer to them for people. To standardize experimental sessions based on personalized sets of stimuli, stimuli were split into three distance categories.

Trials were presented in a randomized block design, with each block containing three consecutive trials belonging to a specific domain and distance category. Each trial was presented for a maximum of 10 seconds (5 TRs). If participants responded faster, trials progressed within two-second windows (if a participant responded 3.5 seconds after trial onset, the trial progressed 4 seconds after trial onset). Each block was followed by 6 seconds of fixation. Participants were instructed to respond accurately but as fast as possible.

In the lexical control task, participants were presented with stimuli pairs from the same sets but were instructed to indicate which of the words contains the letter "A". The lexical control task was administered via two experimental runs, each containing 12 three-trial domain-dedicated blocks in randomized order, balanced for domains.

*fMRI single participant analysis*

A first level general linear model (GLM) analysis was applied to each participant separately. Predictors were constructed for all task conditions and convoluted with a canonical hemodynamic response function. Motion parameters were added to the GLM model to further minimize motion related effects. The constructed model was independently fitted to the time course of each voxel. Analyses were performed for each participant separately in a fixed-effect manner by joining the different experimental runs. This analysis produced task related parameter coefficients (β values) maps.

### Supplementary Results

#### Patterns of neurodegeneration across the AD continuum overlap DMN sub-networks differently

To characterize our sample through well-established neurodegeneration markers of AD pathology, we estimated the differences in GM density and in FDG uptake between AD dementia, MCI, and CN participants across the whole brain and within DMN subnetworks.

VBM analysis showed no significant differences contrasting CN with MCI participants (Figure S1A1). CN and MCI greater than AD dementia participants VBM contrasts revealed a set of overlapping brain regions including the superior middle and inferior temporal gyri, posterior, middle and anterior cingulate cortices, precuneus, parahippocampal gyrus and hippocampus bilaterally (Figure S1A). For GM density in the DMN, significant differences were found only between CN and AD dementia participants in DMN A, B and C (ANOVA and Tukey-Kramer post-hoc test, $P < 0.05$, Figure S2D).

Glucose metabolism was estimated via the use of FDG PET and acquired as complementary metric of neurodegeneration. GLM-derived, CN greater than MCI participants contrast revealed significant hypometabolism in a set of brain regions, including the posterior and middle cingulate cortices, precuneus, parahippocampal gyrus and hippocampus bilaterally. Similarly, MCI greater than AD dementia participants contrast revealed significant hypometabolism in the inferior temporal gyri, posterior cingulate cortex, precuneus, and hippocampus (Figure S1B). The CN greater than AD dementia participants contrast revealed significant hypometabolism in supramarginal, angular, superior middle and inferior temporal gyri, posterior, middle and anterior cingulate cortices, precuneus, parahippocampal gyrus and hippocampus bilaterally (Figure S1B). Significant differences in FDG SUVr were found between CN and AD dementia participants in the brain regions comprising the DMN A subnetwork, and between all groups in DMN C (ANOVA and Tukey-Kramer post hoc test, $P < 0.05$).

**Suplemntary Figures**

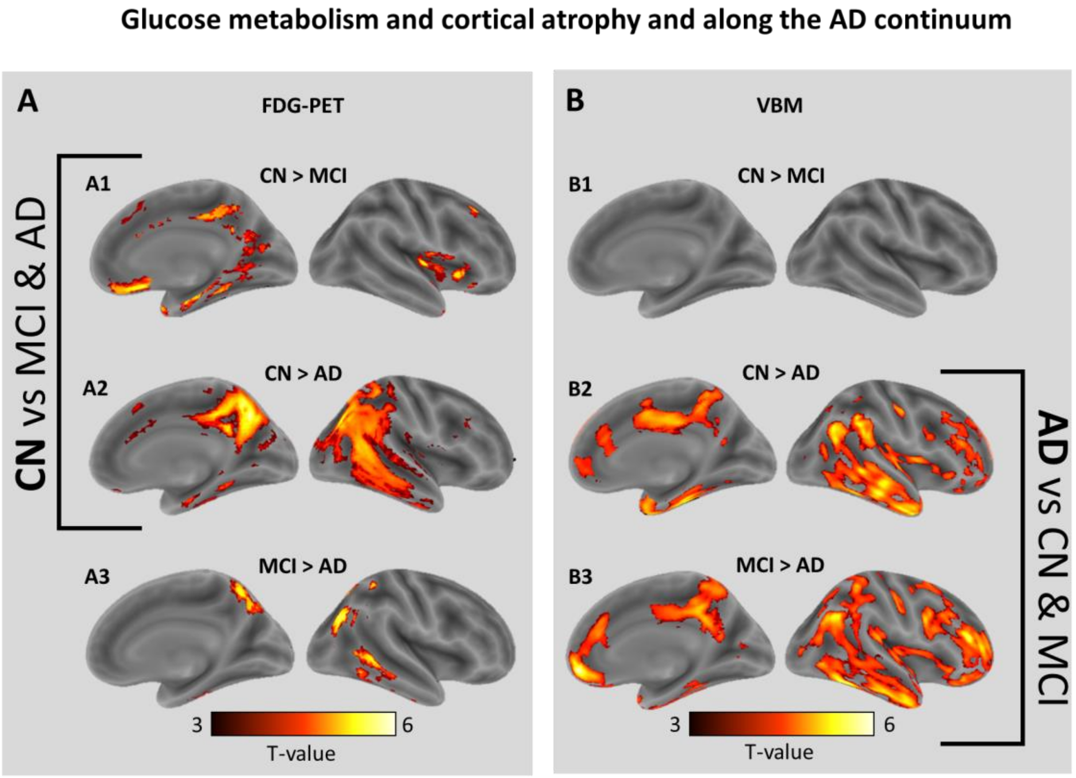


**Figure S1. Glucose metabolism and cortical atrophy along the AD continuum.**

**(A)** Results of GLM analysis, showing projected T-maps of voxels with significantly increased FDG uptake (SUVr values; cerebellar grey matter as reference region) in (A1) CN greater than MCI participants, (A2) CN greater than AD dementia and (A3) MCI greater than AD dementia. **(B)** Results of VBM analysis, showing projected T-maps of voxels with significantly increased GM density in (B1) CN greater than MCI participants, (B2) CN greater than AD dementia and (B3) MCI greater than AD dementia. All voxel surpassed $P<0.05$ (FDR corrected) significance (CN>MCI: DF=38, CN>AD dementia: DF=27, MCI?AD dementia: DF=34), and underwent cluster size thresholding of 20 voxels.

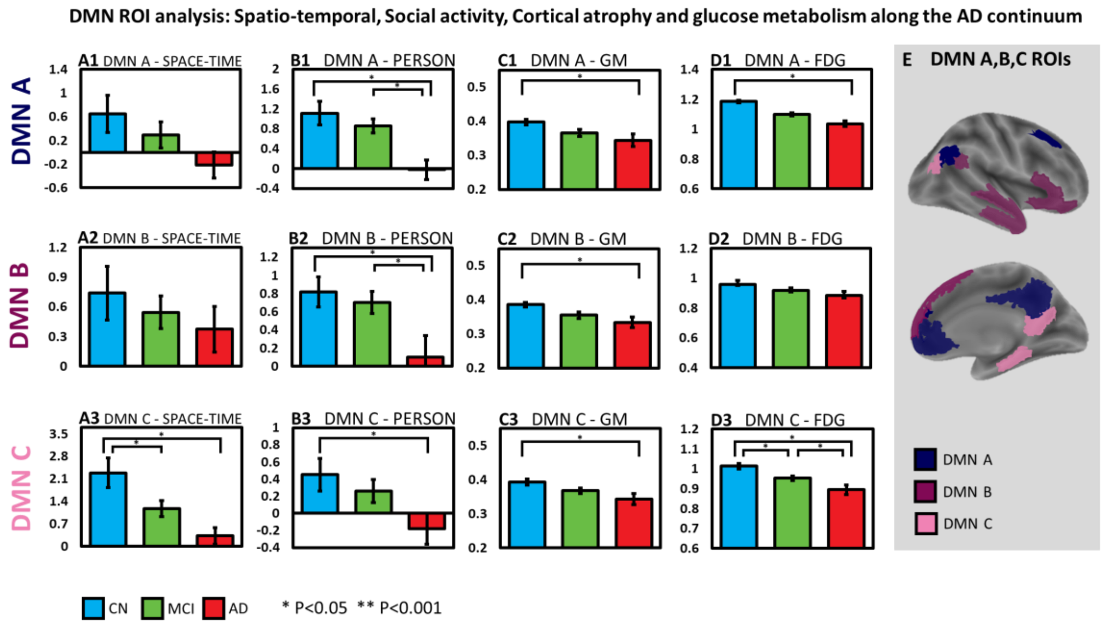


**Figure S2. DMN ROI analysis: Spatio-temporal, Social activity, Cortical atrophy and glucose metabolism along the AD continuum**. Given established DMN association to the AD pathological process, **(E)** DMN A, B, C subnetworks were used as ROIs in which **(A)** spatio-temporal, **(B)** social, **(C)** structural, and **(D)** metabolic mean values were compared between CN (N= 16, blue), MCI (N = 23, green), and AD dementia participants (N = 12, red). **(E)** Segmentation of DMN into DMN A (bright), B (medium) and C (dark) purple clusters was performed, according to the Schaefer parcellation (Schaefer et. al, 2018). For mean parameter estimates of spatio-temproal orientation, significant differences were found between (A3) CN and MCI participants, and CN and AD dementia participants in DMN C (ANOVA and Tukey-Kramer post hoc test, $P < 0.05$). For mean parameter estimates of social orientation (over rest), significant differences were found between (B1-3) CN and AD dementia participants in DMN A, B, and C, and between (B1-2) MCI and AD dementia participants in DMN A and B (ANOVA and Tukey-Kramer post hoc test, $P < 0.05$). For mean GM density values, significant differences were found between (C1-3) CN and AD dementia participants in DMN A, B, and C (ANOVA and Tukey-Kramer post hoc test, $P < 0.05$). For mean FDG SUVr (cerebellum as reference tissue) significant differences were found between CN and AD dementia participants in (D1) DMN A and between (D3) all clinical groups in DMN C (ANOVA and Tukey-Kramer post hoc test, $P < 0.05$).

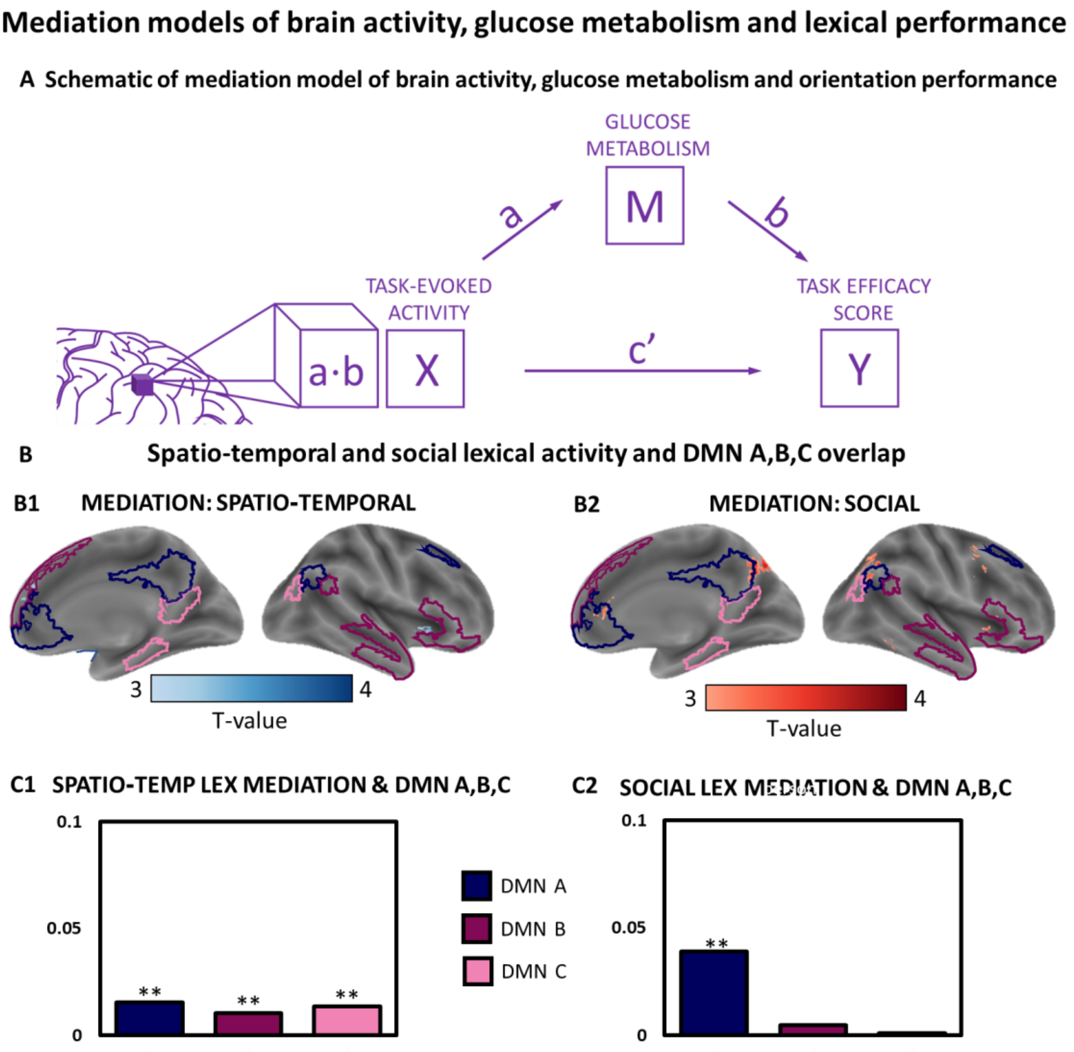


**Figure S3. Mediation analysis of brain activity, glucose metabolism and lexical control performance.** The mediation analysis tested the hypothesis that changes in FDG-PET uptake (M) across the AD spectrum, alters the relations between lexical control-evoked brain activity (X) and lexical control task performance (Y), separately for lexical space-time, and person. For lexical control in space time, mediation effects were found to be significant ($P<0.05$, FDR-corrected) in the insula and the superior medial gyrus, not overlapping with DMN A (A1) or DMN C (A2). For lexical control in person, mediation effects were found to be significant ($P<0.05$, FDR-corrected) in the precuneus and the superior medial gyrus, significantly overlapping with medial ROIs of DMN A (4%, $P<0.001$, B1), not overlapping with DMN C (B2)
